# AN ISOCHORE FRAMEWORK UNDERLIES CHROMATIN ARCHITECTURE

**DOI:** 10.1101/068114

**Authors:** Kamel Jabbari, Giorgio Bernardi

## Abstract

A recent investigation showed the existence of correlations between the architectural features of mammalian interphase chromosomes and the compositional properties of isochores. This result prompted us to compare maps of the Topologically Associating Domains (TADs) and of the Lamina Associated Domains (LADs) with the corresponding isochore maps of mouse and human chromosomes. This approach revealed that: 1) TADs and LADs correspond to isochores, *i.e.,* isochores are the genomic units that underlie chromatin domains; 2) the conservation of TADs and LADs in mammalian genomes is explained by the evolutionary conservation of isochores; 3) chromatin domains corresponding to GC-poor isochores (*e.g.,* LADs) interact with other domains also corresponding to GC-poor isochores even if located far away on the chromosomes; in contrast, chromatin domains corresponding to GC-rich isochores (*e.g.,* TADs) show more localized chromosomal interactions, many of which are inter-chromosomal. In conclusion, this investigation establishes a link between DNA sequences and chromatin architecture, explains the evolutionary conservation of TADs and LADs and provides new information on the spatial distribution of GC-poor/gene-poor and GC-rich/gene-rich chromosomal regions in the interphase nucleus.

---

By assessing the probability of physical proximity between pairs of loci, the chromosome conformation capture (3C) approach (Dekker et al. 2002) and its further developments, such as Hi-C (Lieberman-Aiden et al. 2009) and *in situ* Hi-C (Rao et al. 2014), have revolutionized our understanding of mammalian chromatin architecture. The three-dimensional structure of chromatin is now seen as consisting of topologically associating domains (TADs), a system of chromatin loops and boundaries 0.2-2 Mb in size (Dixon et al. 2012; Nora et al. 2012), the majority of which can be resolved into contact domains, 185 Kb in median size (Rao et al. 2012). Another approach, DamID technology, has shown that lamina-associated domains (LADs) have a median size of 500 Kb are GC-poor and are scattered over all chromosomes (Meuleman et al. 2013; Kind et al. 2015). It is also well established that TADs and contact domains are largely conserved across different mouse and human cell types (Dixon et al. 2012; Rao et al. 2014; Vietri Rudan et al. 2015). Moreover: 1) the regulation of gene expression is affected by topological changes such as boundary deletions and domain perturbations (see Ghirlando and Felsenfeld, 2016; Merkenschlager and Nora, 2016; Dixon et al. 2016; Pueschel et al. 2016; Gonzalez-Savon and Gasser, 2016; Sati and Cavalli, 2016; Solovei et al. 2016, for recent reviews); and 2) senescence is accompanied by alterations in LAD/lamina interactions (Chandra et al. 2015). However, the formation mechanism of topological domains is not yet understood, in spite of the recent advances in the field of chromatin architecture and of the interesting models proposed so far (Lieberman-Aiden et al. 2009; Rao et al. 2014; Vietri Rudan et al. 2015; Sanborn et al. 2015; Dekker and Mirny, 2016; Rowley and Corces, 2016; see Dixon et al. 2016, for a critical review).

The investigations just mentioned (and many others not cited here) were mainly focused on the connections between interphase chromatin structure and gene expression/regulation and replication on the one hand and on chromatin/lamina interactions on the other. We were aware for a long time, however, of the existence of correlations between the base composition (and related properties) of isochores and the structural features of chromosomes (see Supplemental Tables S1-S2 and Bernardi, 2015). Moreover, a recent analysis revealed that correlations exist between the two main compositional compartments of isochores (the large GC-poor/gene-poor genome desert and the small GC-rich/gene-rich genome core) and the architecture of chromatin (see Supplemental Table S3 and Bernardi, 2015). These correlations encouraged us to investigate in detail the genomic basis of chromatin architecture. This was done by comparing maps of TADs and LADs with the corresponding maps of isochores from mouse and human chromosomes and by looking for an explanation for the evolutionary conservation of TADs.

## TADs, LADs and isochores from human and mouse chromosomes

The isochores/TADs comparison of mouse chromosomes indicated a good degree of match between the boundaries of TADs and the boundaries of the corresponding isochores or short isochore blocks, as outlined in Fig. 1A for chromosome 17. In the case of GC-poor isochores, the correspondence not only concerned the self-interactions and the short-range interaction frequencies on the diagonal of the contact frequency matrix, but also the long-range ones along the two major axes. The latter interactions were found to concern domains that correspond to other, even faraway, GC-poor isochores. In contrast, TADs corresponding to GC-rich isochores showed strong interactions only with nearby TADs also corresponding to GC-rich isochores. At a higher resolution (100 Kb *vs*. 300 Kb), boundaries corresponding to isochore blocks (such as the rightmost one) appeared to be often resolved into boundaries corresponding to individual isochores (see Fig. 1B).

**Fig. 1.**
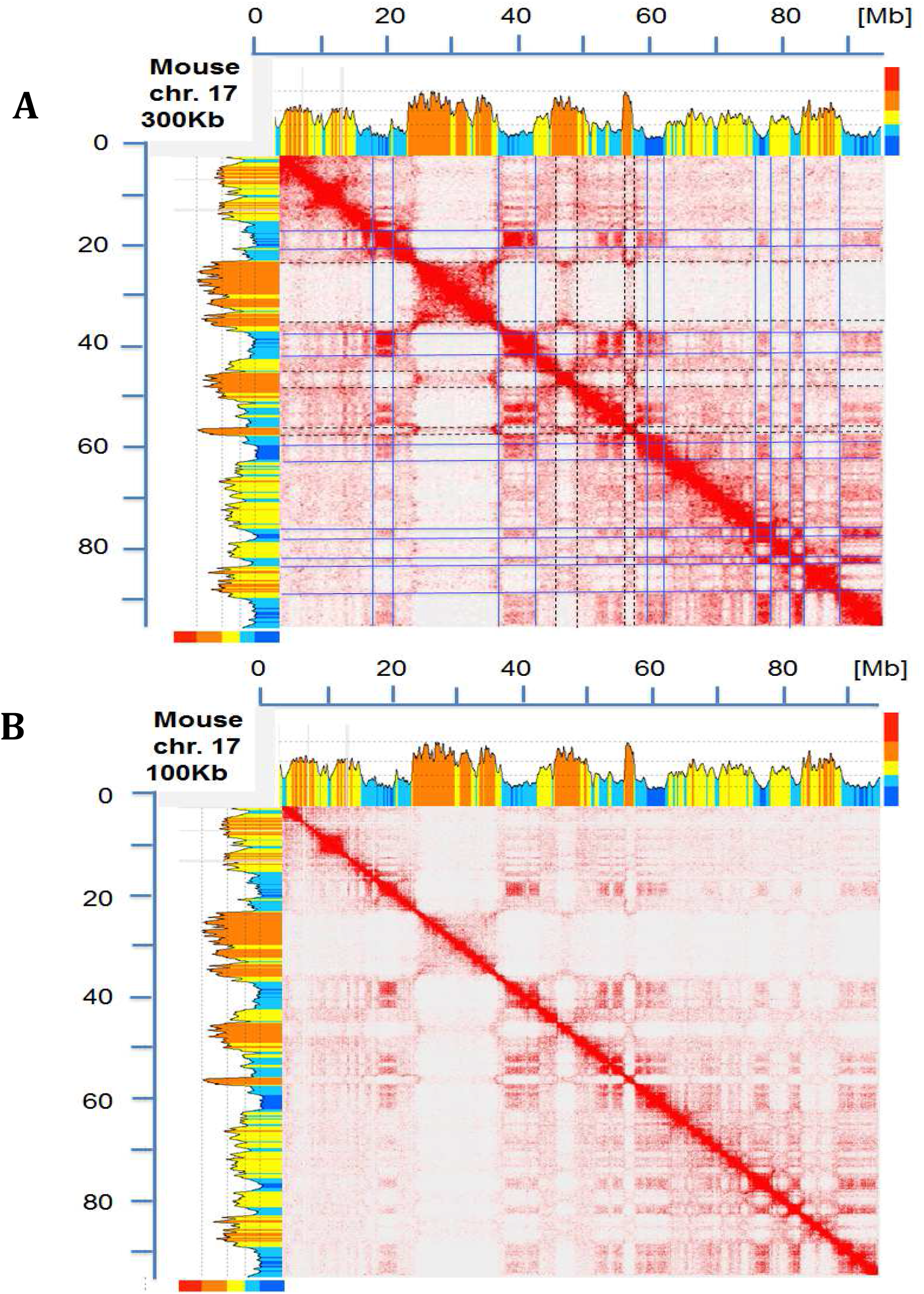
Figure 1A. The heat map of chromatin interactions in mouse chromosome 17 (from Vietri Rudan et al. 2015) is compared with the corresponding compositional profile (drawn from mm 10 genome assembly using a sliding window of 300 Kb and the program of Paces et al. 2004). Isochore families L1 to H3, characterized by increasing GC levels are defined according to the “fixed” boundaries between isochore families (see Supplemental Table S1) and are represented in different colors, deep blue, light blue, yellow, orange and red, respectively; the multicolored vertical bars on the top right indicate GC levels that correspond to the compositional boundaries among isochore families. The self-interactions along the diagonal, as well as the interactions along the two major axes, correspond to short isochore blocks or to individual isochores, as stressed by lines through the coinciding boundaries of TADs (blue and broken black lines correspond to GC-poor and GC-rich isochores or isochore blocks, respectively; not all lines were drawn to avoid readability problems). Interactions corresponding to GC-poor regions are also seen in domains corresponding to other GC-poor regions even when located far away on the chromosomes, the intensity of such interactions decreasing, however, with distance. In the case of interactions corresponding to GC-rich isochores, much weaker signals are present outside the diagonal, an indication of more localized interactions. Note that mouse chromosome 17 is an acrocentric chromosome and that the centromeric sequences correspond to the region near the origin of the Megabase (Mb) scale. Unless otherwise stated, all the interaction maps presented in this article are shown at a 250 Kb resolution and isochores are visualized using a 300 Kb sliding window across chromosomes. Figure 1B. The heat map of mouse chromosome 17 is compared at a resolution of 100 Kb with the corresponding isochore profile. Interactions along the diagonal are weaker because split into smaller domains, that reveal fine details. For example, the largest GC-rich region on the chromosome (around 30 Mb) is now resolved into several domains that present a finer correspondence with isochores.

Results for human chromosomes (see Figs. 2A and 2B for chromosome 7 at resolutions of 300 Kb and 50 Kb, respectively) were similar to those of Fig. 1 (see the rightmost block of Fig. 2B) in spite of the greater compositional complexity of the human genome (in which isochores extend further on both GC-poor and GC-rich sides and cover a ~25% GC range; Bernardi, 2015).

**Fig. 2.**
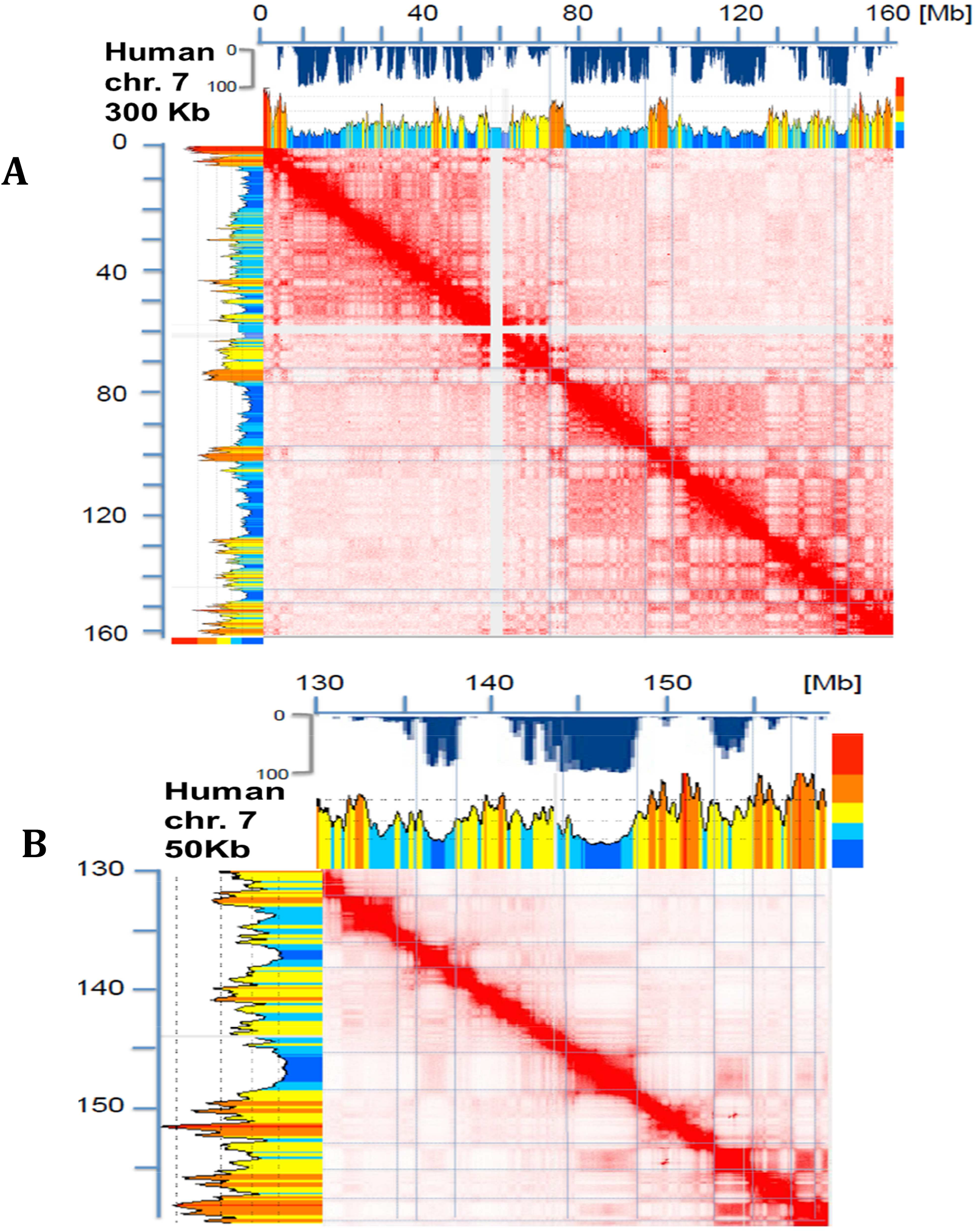
Figure 2A. The heat map of chromatin interactions of human chromosome 7 (from Rao et al. 2014) and the corresponding LAD map (from Kind et al. 2015; blue inverted profile under the Megabase axis) are compared with the corresponding compositional profile (using the matched b37 assembly of Rao et al. 2014, and the program of Paces et al. 2004). The results are comparable with those of Fig. 1A, except that the centromeric repeats (that have been sequenced in this chromosome) do not interact with any other sequence and are responsible for the two orthogonal blank stripes on the heat map. Blue lines correspond to TADs, LADs and inter-LADs. Figure 2B. The heat map of chromatin interactions over ~30 Mb of human chromosome 7 is analyzed at a resolution of 50 Kb. In this case a finer correspondence of isochore boundaries with LAD and TAD boundaries can be observed.

In this case, LAD maps were also available (Kind et al. 2015) and could be compared with isochore maps. This revealed a very good match between LADs and GC-poor isochores (as expected from previous work; Meuleman et al. 2013; Kind et al. 2015) and no match between LADs and GC-rich isochores. The findings of Figs. 1 and 2 (that were extended to all mouse and human chromosomes; see a later section) indicate that TADs and LADs appear to correspond to individual isochores. Indeed, the cases in which TAD boundaries correspond to short isochore blocks seem to be essentially due to the structural complexity of a number of TADs, to the comparison of TADs from a given source with a genomic sequence from a different source (see legend of Fig. 2A), and also to some oversegmentation of isochores due to the use of “fixed” boundaries” for the definition of isochore families (see Supplemental Table S1).

The correspondence of TADs and LADs with isochores is also supported 1) by the independent evidence that isochores are replication units, characterized by all early or all late replicons in GC-rich and GC-poor isochores, respectively (Costantini and Bernardi, 2008), and that TADs are stable units of replication timing (Pope et al. 2014); 2) by the fact that the ~2,200 TADs (Dixon et al. 2012) and 1,300 LADs (Kind et al. 2015) together approximately match the 3,200-3,400 isochores of the human genome (Bernardi, 2015); 3) by the match of the size range of TADs (0.2-2 Mb; see Dixon et al. 2012; Nora et al. 2012) and LADs (~0.5Mb) with that of isochores (0.9 Mb, number average; Costantini et al. 2006), and of the larger average size of LADs compared to TADs with the larger size of GC-poor compared to GC-rich isochores (Costantini et al. 2006).

The information which is available on isochore families (see Supplemental Table S1), on TADs and LADs also is of interest in connection with the composition of the architectural elements of chromatin: 1) ~35% of the genome consist of GC-poor sequences that correspond to LADs (Meuleman et al. 2013) and comprise all the L1 isochores (that represent 19% of the genome) plus ~ 16% of the genome corresponding to L2 isochores; 2) since the total amount of GC-poor isochores, from the L1, L2 and H1 families, represents 86% of the genome, then 86%-35%, *i.e.,* ~50% of the genome consists of GC-poor sequences not present in LADs, but present in TADs; since the latter also include 14% of the genome made by GC-rich sequences from H2 and H3 families, then ~22% (14/64) of TADs are GC-rich and ~78% are GC-poor; 3) the properties of chromatin boundaries match those of GC-rich isochores; in fact, boundaries should be visualized as short sequences that are GC-richer than loops and share properties with GC-rich isochores (see Supplemental Table S3).

## Higher resolution comparisons of TADs and isochores

Fig. 3 presents a series of chromatin loops (that can be identified with TADs) from a 2.1 Mb segment of human chromosome 20 (data from Rao et al. 2014) along with the corresponding contact matrix (also from Rao et al. 2014). Red and blue horizontal broken lines connect “loop domains” (anchored by CTCF, the CCCTC-binding factor) and “ordinary domains” (not anchored by CTCF), respectively, with the corresponding squares in the heat map. Thin vertical lines along the borders of the squares (double lines corresponding to intervals between loops whenever large enough) were used to segment the corresponding compositional profile of DNA. This approach was shown to segment DNA into a series of alternating H1 and H2 isochores and to establish a correspondence between chromatin loops and isochores (which is stressed by the square brackets under the compositional profile).

**Figure 3.**
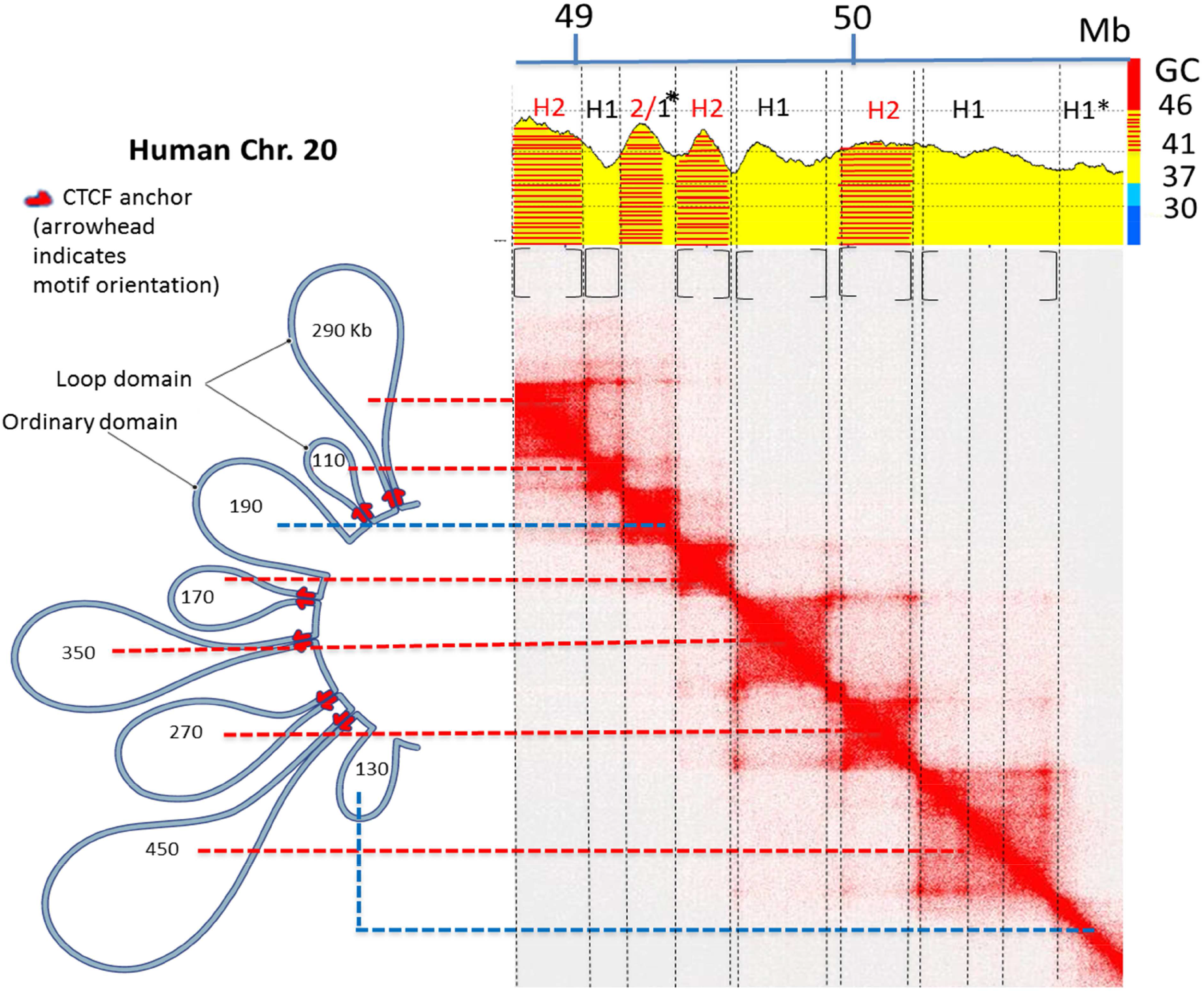
The chromatin loops from a 2.1 Mb region of human chromosome 20 (Fig. 6F from Rao et al. 2014) have been aligned with the corresponding heat map which was used to segment the corresponding DNA sequence into isochores. In this Figure the “extended” isochore ranges of Supplemental Table S1 were used to assign isochores to families, in order to take care of some minimal trespassings of the 46% GC upper threshold of H1 isochores. Asterisks indicate anomalies in the isochores/domains correspondence (see Text).

It should be noted 1) that the only exceptions to the alternating isochore rule concerns the two “ordinary domains”: the 190 Kb domain encompasses a H2/H1 sequence and the 130 Kb domain is formed by a H1 isochore which follows another H1 isochore; 2) that the largest 450 Kb loop consists of three sub-domains that are clearly visible on the heat map; and 3) that the 2.1 Mb region does not present any LAD signal, as expected from its GC-richness.

## Additional observations on isochores, TADs and LADs

A number of other observations are relevant: 1) the density of CTCF binding sites is decreasing in isochores from families of increasingly lower GC and especially low in family L1 (Supplemental Fig. S1A); 2) inter-chromosomal interactions (from Rao et al. 2014) have a very strong preference for GC-rich regions (see Fig. 4); this point, as well as the previous one, will be commented in the Conclusions; 3) a comparison of isochore profiles with the sub-compartments of contact domains (from Rao et al. 2014) shows that, in general, A1 sub-compartments correspond to H2/H3 isochores (sometimes including flanking isochores from the H1 and even from the L2 family), A2 sub-compartments to H1 and L2 isochores, B1-B3 sub-compartments to L2 and L1 isochores (see Supplemental Fig. S1B and its legend for further comments); needless to say, these results provide a more detailed picture (in the direction of isochore families) of the previously noted general correspondence (Bernardi, 2015) between (i) the open A and closed B chromatin compartments (Lieberman-Aiden et al. 2009) and (ii) the genome core and the genome desert; 4) Supplemental Figures S2-S20 and S21-S42 display the correspondence of isochores with TADs (and LADs) along all mouse and human chromosomes, respectively, so demonstrating the generality of the results of Figs. 1 and 2.

**Figure 4.**
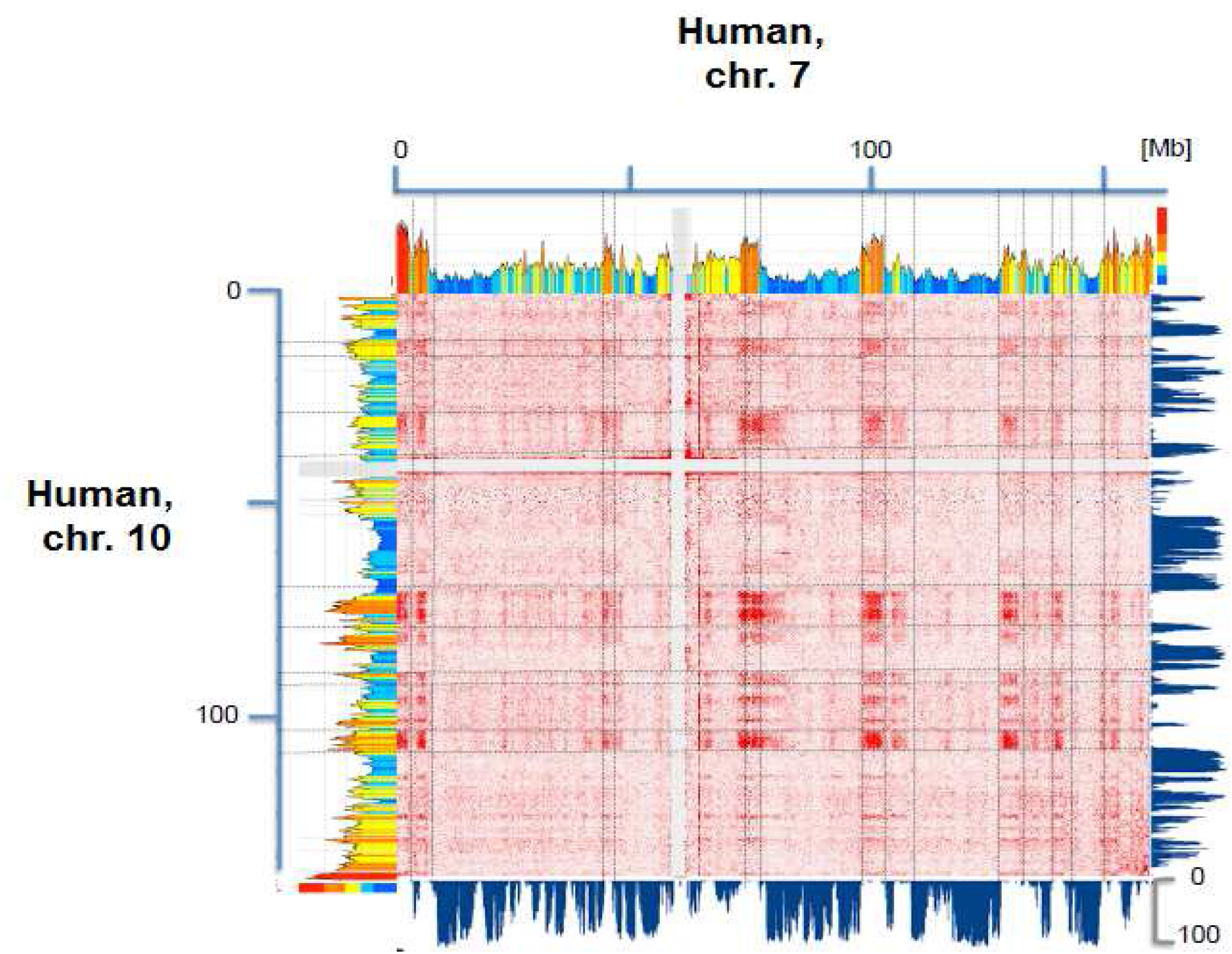
The heat maps of chromatin interactions of human chromosomes 7 and 10 (from Rao et al. 2014) are compared with the corresponding compositional and LAD profiles (from Kind et al. 2015). Interactions appear to correspond to GC-rich isochores and inter-LADS and to be widely spread over the two chromosomes. Lines are used to guide the visual inspection of these features.

A detailed statistical analysis of isochores and TADs (from Dixon et al. 2012) based on permutation tests (Gel et al. 2015) showed a non-random associations with a *p*-value <0.001; moreover, it was demonstrated that the association between isochores and TADs was specifically linked to common boundary positions (see Supplemental Figs. S43-S44 and comments in Supplemental Material).

## Evolutionary conservations of TADs and isochores

As far as the evolutionary conservation of TADs is concerned, we examined the possible connections between the conservation of isochores (Pavlicek et al. 2002; Bernardi, 2007) and the conservation of chromatin domains (Rao et al. 2014; Dixon et al. 2012; Vietri Rudan et al. 2015) in syntenic loci. Fig. 5 compares two syntenic regions of human chromosome 14 and mouse chromosome 12 and shows that the isochore profiles are almost mirror images of each other (Fig. 5A) and that the regions correspond to similar heat maps of chromatin interactions (Fig. 5B). Figs. 5C and 5D show the correlations between GC levels of human isochores and the GC levels of 1,348 and 1,358 syntenic regions from mouse and dog, respectively. In both plots, highly significant correlations were found across the whole GC spectrum. This is an indication that GC-poor isochores (many of which correspond to LADs) are also conserved in evolution, in agreement with the evolutionary conservation of isochore families (Costantini et al. 2009). This provides an additional point in favor of the correspondence of isochores with TADs and LADs, because the conservation of isochores accounts for the conservation of chromatin architecture.

**Fig. 5.**
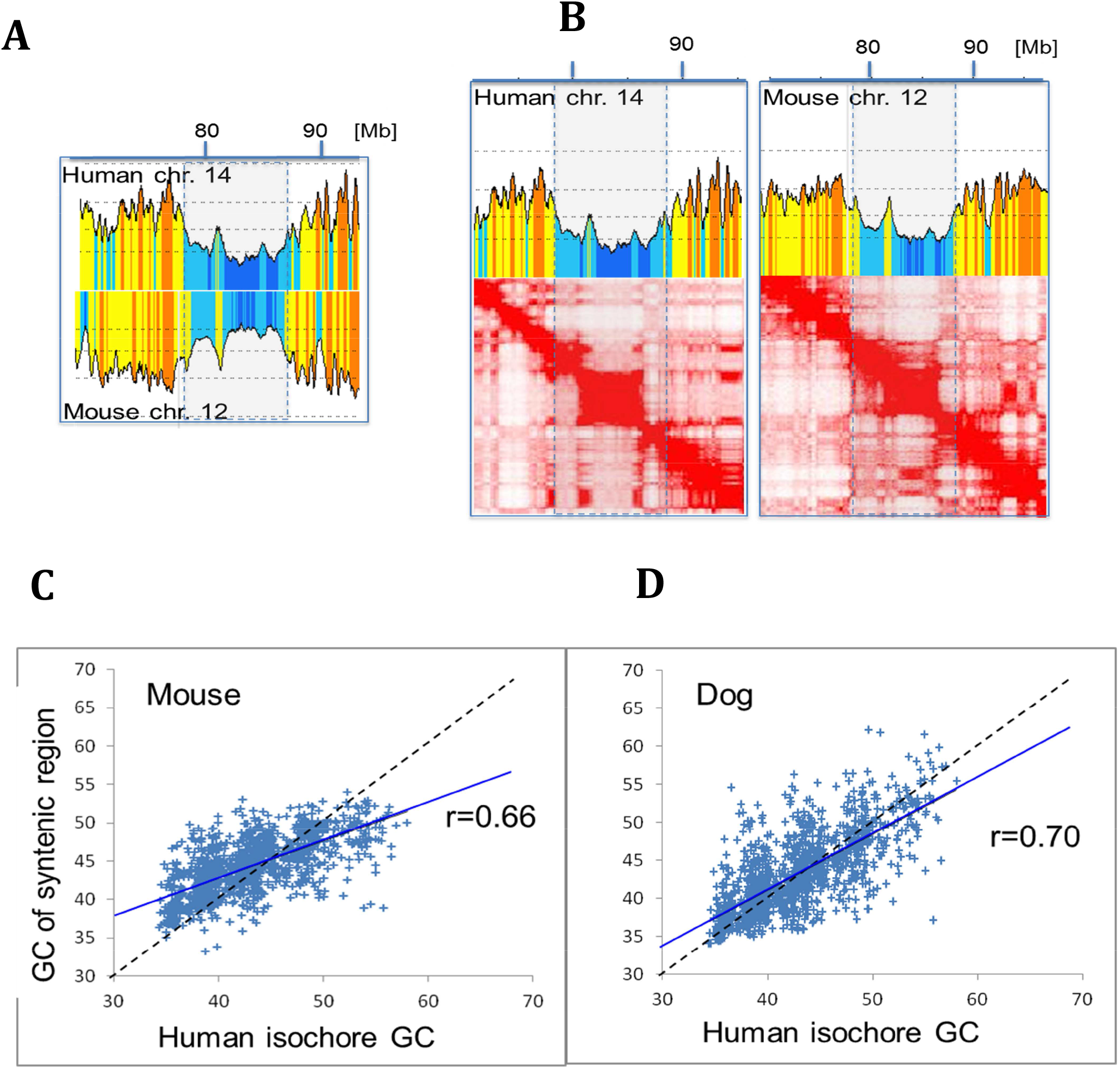
Figure 5A. The compositional profiles of two syntenic regions located on human chromosome 14 and mouse chromosome 12 (from Rao et al. 2014 and Vietri Rudan et al. 2015, respectively) are almost mirror images of each other, showing the compositional conservation of syntenic regions in evolution. Figure 5B. A comparison of compositional profiles and heat maps (from Rao et al. 2014) of the two syntenic regions from Fig. 5A. Figure 5C,D. GC levels of 1,348 and 1,358 syntenic regions from mouse (left) and dog (right) are plotted against the GC levels of human isochores.

## Conclusions

The main conclusions of the present investigations are presented below.

1) The long-distance interactions of GC-poor domains mainly concerning LADs, on the one hand, and the “local” and interchromosomal interactions of GC-rich domains, on the other hand, provide a view of chromosomes in the interphase nucleus which is in line with results obtained using different approaches. Indeed, previous results indicated that GC-rich, early replicating, transcriptionally active chromatin regions are located in the nuclear interior (Sadoni et al. 1999) and that the gene-rich regions display a much more spread-out conformation (Saccone et al. 2002; Gilbert et al. 2004). We understand now that this facilitates interchromosomal interactions that are not possible in the case of the compact gene-poor regions located at the nuclear periphery. In contrast, LADs (that correspond to GC-poor isochores) exhibit long range interactions that are, however, linked to the same chromosomes, a strong indication that such LADs may share a remarkable degree of contiguity on the lamina.

2) The isochore/loop correspondence implies a fair compositional homogeneity (a typical isochore property), as well as the existence of different GC levels and different frequencies of di- and tri-nucleotides (Costantini and Bernardi, 2008b), in different TADs and LADs (as shown in Fig. 3 and in Supplemental Figs. S2-S42); in other words, while the genome is a compositional mosaic of isochores (Bernardi et al. 1985; Bernardi, 2004), interphase chromatin is a mosaic of different TADs and LADs. In turn, this is in agreement with different structural/functional properties of TADs (see Supplemental Tables S2 and S3), whereas in the case of LADs, L1 isochores possibly correspond to the “stable contacts”, L2 isochores to the “variable contacts” of Kind et al. (2015; see also the legend of Supplemental Figs. S21-S42).

3) The key finding of these investigations, namely the correspondence of TADs and LADs with isochores, indicates that isochores should be visualized as the framework of chromatin architecture. In fact, the present results lead to a new paradigm of chromatin architecture which was obtained by taking into consideration its DNA backbone, the isochores, “a fundamental level of genome organization” (Eyre-Walker and Hurst, 2001), and by showing that isochores are the genomic units that underlie TADs and LADs. This establishes a connection between DNA sequences and chromatin architecture and opens the way to studying changes in chromatin architecture at the sequence level.

## Acknowledgments

The authors are indebted to Giacomo Bernardi and Oliver Clay for critical reading, detailed comments and discussions, and to Caterina Nuvoli for excellent technical help.

